# Calibrating Hall-Effect valvometers accounting for electromagnetic properties of the sensor and dynamic geometry of the bivalves shell

**DOI:** 10.1101/2020.12.20.423648

**Authors:** Jean-Marc Guarini, Jennifer Coston-Guarini, Luc A. Comeau

## Abstract

Hall-Effect valvometry (HES) is being used to describe bivalve valve gape variations and infer environmental perturbations in a variety of aquatic environments. Surprisingly, the published calibrations in ecological literature ignore both the electromagnetic properties of HES and that the valves rotate around their hinge when they move. The high sensitivity of HES suggests these features should be accounted for explicitly to estimate measurement accurately. To address these issues, two calibration functions were developed based on the electromagnetic properties of the HES: one assumes that the HES and magnet are maintained on the same linear axis, and the second model accounts for the geometric properties of the system (i.e. variations of the angle between HES and the magnet during shell rotation). The great scallop (*Pecten maximus*) was used as biological model because of its large range of valve openings. HES were installed on the flat valve and magnets installed on the opposing rounded valve; 12 individuals of similar size (10 ± 1(SD) cm), were equipped and placed in controlled experimental conditions. A calibration was done for each individual once time series recordings were completed. The variability of parameter estimates was calculated with a bootstrap method. The second model (with rotatation) improves valve gape distance estimates for larger openings despite the decrease of sensor sensitivity. To infer valve gape dynamics, the reciprocal calculation of the calibration function was formalized and applied to the Hall voltage time series. Our analysis suggests that under controlled laboratory conditions, scallops are partially open most of the time (inter-valve distance equal *ca*. 27 mm on average, or 45 % of the average maximum opening distance). Interspersed in this continuous regime, individual scallops performed closing events at a frequency of *ca*. 2.5 closings per hour. A closing event is a movement that is fast enough relative to the recording frequency (10 Hz) to qualify as discrete. We find that the inversed calibration model without rotation allows negative value estimates, which indicates that this calibration function is incorrect. In contrast, the inversed calibration model with valve rotation around the hinge constrains gape distance values in their domain of definition which automatically excludes sensor readings that produce negative values from estimated gape time series.

## 1 Introduction

Bivalve species are widely harvested or farmed and commercialised in coastal zones and these fisheries are often in conflict with other usages. Adults are sedentary or have little mobility. Therefore considerable scientific investigations have been performed to understand how these organisms are affected by local environmental disturbances. Since the 1970s bivalves have been used as sentinel species for coastal ecosystems – mainly for the detection of water-borne contaminants (*e.g*. Goldberg, 1975). At present, diverse bivalve-based monitoring programs exist in many countries (Guégen et al., 2011), for shellfish aquaculture or environmental impact assessment purposes (*e.g*. Steinhardt et al., 2016).

To reinforce monitoring programmes, and converge toward integrated warning systems, one current objective is to obtain ”real-time” responses of living organisms to environmental disturbances *in situ*. One technical mean to achieve this goal is ”valvometry”, or the measurement of valve gape variations, which has been developed first for physiological studies (Marceau, 1909) and then was expanded to investigate individuals’ behavior as a function of changes of their proximal environment (Gaine and Shumway, 1988; Nagai et al., 2006; Redmond et al., 2017). Valve closures result from a contraction of the adductor muscle, which opposes a continuous opening force generated by the ligament at the umbo (Trueman, 1953). Therefore, valve movements depend on metabolism and have a functional and protective roles in bivalves physiology; particularly, environmental stresses can cause voluntary closing of the shell, while increasing metabolic cost in term of energy. Valvometry is today a generic term encompassing a wide range of techniques: from early wire-myographs connected to sooted glass recorders (Marceau, 1909), to automated high frequency sensor-recorder systems, like strain gauges (Wilkens, 1981), Hall-effect sensors (Nagai et al., 2006), impedance electrodes (Tran et al., 2003), and fiber optics (Franck et al., 2007).

Patterns of valve movements have been directly linked to basic biological activities such as respiration, excretion, or feeding (Nagai et al., 2006), and to some behaviours including swimming, spawning, or burying (Thomas and Gruffydd, 1971; Wilkens, 1981). In addition, there is a rich literature describing how variations of valve opening amplitudes and movement frequencies were attributed to changes in temperature in *Crassostrea virginica* (Comeau et al., 2012), salinity in *Mytilus edulis* (Bamber, 2018), presence of toxic algae in *Pinctada fucata* (Nagai et al., 2006) and *Mytilus galloprovincialis* (Comeau et al., 2019), cadmium chloride concentrations in *Corbicula japonica* (Moroishi et al., 2009), contaminants of free metals (*e.g*., copper) in *Corbicula fluminea* (Liu et al., 2016), alkaline solutions (*e.g*., biofouling compounds) in *Mytilus edulis* (Comeau et al., 2017), suspended particulate matter concentrations in *Pecten maximus*, (Szostek et al., 2013), pH changes in *Arctica islandica* (Bamber and Westerlund, 2016), and to noise in *Magallana gigas* (Charifi et al., 2018).

In this article, we focus on Hall-Effect sensors (HES). HES measure variations in the Hall-voltage generated proportionally to the magnetic field strength passing through them (Hall, 1879; Propovic, 2003). These sensors have been used in many engineering applications (Popovic, 2003) and have been tested with bivalves (Wilson et al., 2005; Nagai et al., 2006; Clements et al., 2018), including with *Pecten maximus* (Robson et al., 2009; Szostek et al., 2013). They are small, lightweight, and less invasive than other valvometry systems, as well as being easy to set up, highly sensitive and cost-effective. The HES output (i.e. the Hall voltage) is influenced by the composition of the magnet, the nature of the sensor semi-conductor (Popovic, 2003; Ramsden, 2011) and the environmental conditions (Liebsch, 2002; Ejsing, 2006); the Hall voltage depends on the strength of the magnetic field and the relative distance between the sensor and the magnet. For valvometry applications, this means that the Hall voltage measured will depend not only on the distance between the sensor and magnet but also on their relative positions on individual organisms, as well as valve rotations made during movements. Hence, both the physical properties of the sensor installed, as well as changes in alignment between HES and the magnet should be accounted for during measurement calibration. Unexpectedly, in our review of the literature, we found that this has not yet been done. In addition, since monitoring programs are often operating in politically sensitive contexts, this oversight in the calibration methodology could become problematic.

We address explicitly these calibration issues and propose the means to estimate valve gape distances and their uncertainty from time series of Hall-voltage measurements. For this, we designed a calibration function that accounts for the electro-magnetic principles of HES sensors. Then, two calibration models were developed: a first one that assumes the HES and magnet are maintained on the same linear axis (*i.e*. a simple translation of HES relative to the magnet position), and a second one that accounts for the geometric properties of the system (*i.e*. variations of the angle between HES and the magnet during shell rotation around the hinge axis). We then inversed these two calibration functions to calculate valve gape distances from time series of HES output under experimental conditions. To test our calibration method, we used measurements performed on great scallop *Pecten maximus* individuals. this species present a strong interest in monitoring stock conditions, because of its high economic value (Shumway and Parsons, 2016), but it was chosen mainly because individuals exhibit large valve gape opening amplitudes, hence maximize rotation effects on valve gape estimates. We evaluated the resulting patterns in valve gape dynamics relative to described behaviours (Baird and Gibson, 1956; Thomas and Gruffydd, 1971; Wilkens, 1981; Brand, 1991; Robson et al., 2012; Shumway and Parsons, 2016), and compared properties of the dynamic patterns infered from estimates from the two inverse functions.

## 2 Concepts and Methods

### 2.1 Calibration models to quantify the relationship between inter-valve distances and sensor readings

HES Valvometry sensors operate under the Hall-effect principle (Hall, 1879). When an electric current is induced in the HES (Figure 1B), it responds with an output voltage (the Hall effect voltage, *V_H_*) proportional to the orthogonal component of the magnetic field strength applied to it. *V_H_* is therefore a function of the intensity of the induced electric current *I_E_* and the magnetic field strength, *B* (Popovic, 2003) (Equation 1, Table 1). In the ideal case, the proportionality is ensured by the Hall constant, *K_H_*, which is a function of the composition of the sensor and environmental conditions (Kadri et al., 1984).

**Figure 1:**
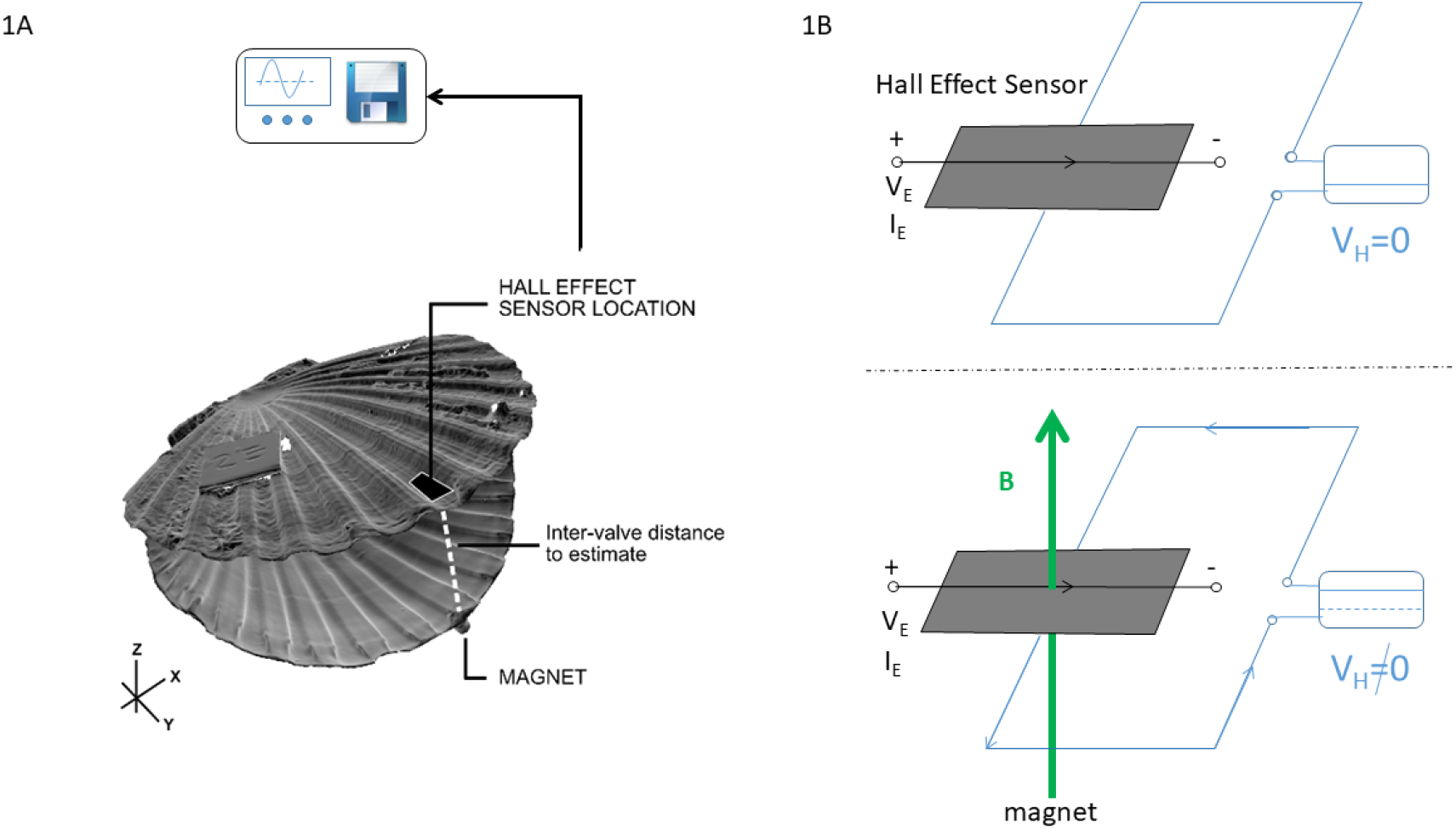
Panels: (A) Schematic of a scallop (*Pecten maximus*) shell showing the placement of the Hall-effect sensor (HES) on the flat valve and the magnet on the curved valve and (B) a schematic depicting the functioning of an HES without being submitted to a magnetic field (upper figure) and while being submitted to a magnetic field (green arrow, lower figure). The HES is powered by an electric current with voltage *V_E_* and intensity *I_E_*, but does not produce any current if not submitted to a magnetic field. When submitted to a magnetic field, the HES produces a current perpendicular to the current *V_E_, I_E_* imposed upon it. Then a Hall-effect voltage (*V_H_*) can be measured, which is proportional to the inverse square distance between the sensor and the magnet.

**Table 1.** Formulations used to estimate the inter-valve distance from valvometry measurements with Hall-effect sensor. Numbers (Nb) are referenced in this article. M.1 and M.2 correspond to Model 1 and Model 2 respectively.

| Nb | Formulation | Description |
| --- | --- | --- |
| 1 | $V_H(t) \propto B(t)I_E$ | $V_H(t)$ (volt): Hall voltage<br>$B(t)$ (tesla): Magnetic field<br>$I_E$ (ampere): Current intensity |
| 2 | $B(t) \propto 1/((D_e(t))^2 \cos(\kappa(t)))$ | $D_e(t)$ (mm): dist. HES-Magnet<br>$\kappa(t)$ (radian): angle between the axis of the magnetic field and axis between the HES and the magnet faces |
| 3 | $D_m(t) = d(t)R/L$ | $D_m(t)$: Measured dist. (mm)<br>$L$ (mm): dist. Umbo-HES<br>$R$ (mm): dist. Umbo-Edge |
| **M.1 Without rotation ($\kappa(t) = 0$)** |  |  |
| 4 | $D_e(t) = d(t) + d_0$ | $d_0$ (mm): fixed dist.<br>$d(t)$ (mm): Inter-valve dist. |
| 5 | $U_H(d(t)) = \theta_1/(d(t) + \theta_2)^2$ | Calibration Function<br>$U_H$ (unitless): HES output<br>$\{\theta_1, \theta_2\}$: parameters to estimate |
| 6 | $d(t) = (\theta_1/U_H(d(t)))^{0.5} - \theta_2$ | Inversed Calibration Function |
| **M.2 With rotation ($\kappa(t) \in ]0, \pi/2[$)** |  |  |
| 7 | $D_e(t)^2 = d_0^2 + d(t)^2 - 2d_0d(t)\cos(\beta(t) + \pi/2)$<br>$\beta(t) = \cos^{-1}(d(t)/(2L))$ | $d_0$ (mm): fixed dist.<br>$d(t)$ (mm): Inter-valve dist. |
| 8 | $U_H(d(t)) = (\theta_1/(A_1^2 \cos(\kappa(t)))$<br>$A_1 = \theta_2^2 + d(t)^2 - 2\theta_2d(t)\cos(\beta(t) + \pi/2)$<br>$\kappa(t) = \cos^{-1}(\frac{D_e(t)^2 - d(t)^2 + \theta_2^2}{2D_e(t)\theta_2^2})$ | Calibration Function<br>$U_H$ (unitless): HES output<br>$\{\theta_1, \theta_2\}$: parameters to estimate |
| 9 | $A_2\sqrt{A_3} = (\theta_1/U_H(d(t)))$<br>$A_2 = \theta_2 + d(t)\sin(\cos^{-1}(d(t)/(2L)))$<br>$A_3 = d(t)^2 + \theta_2^2 + 2d(t)\theta_2\sin(\cos^{-1}(d(t)/(2L)))$ | Inversed Calibration Function |
| **Criteria function** |  |  |
| 10 | $y = \sum_{i=1}^{16} (w_i(U_H(d_i) - U_{H_i})^2)$<br>$w_i = d_i / \sum_{i=1}^{16} d_i$ | Weighted least squares criterion |

The magnetic field strength B varies as a function of the inverse of the square distance, *D_e_* (in meters), between the magnet and the HES (Figure 2A). It depends on how the sensor and magnet are positioned relative to one other. Introducing *κ*(*t*), the angle between the axis of the magnetic field and the axis between the HES (Figure 2A) and the magnet face, the magnetic field strength was assumed to be expressed as Equation 2 (Table 1). When *κ*(*t*) is equal to zero, the HES is assumed to be aligned with the magnet face (i.e., they are in two parallel planes and *cos*(*κ*(*t*)) is equal to 1.0). The limit of functioning of the system is reached when *cos*(*κ*(*t*)) is equal to 0.0 (*κ*(*t*) = 0.5*π*), hence when the sensor is positioned orthogonally from the magnet.

**Figure 2:**
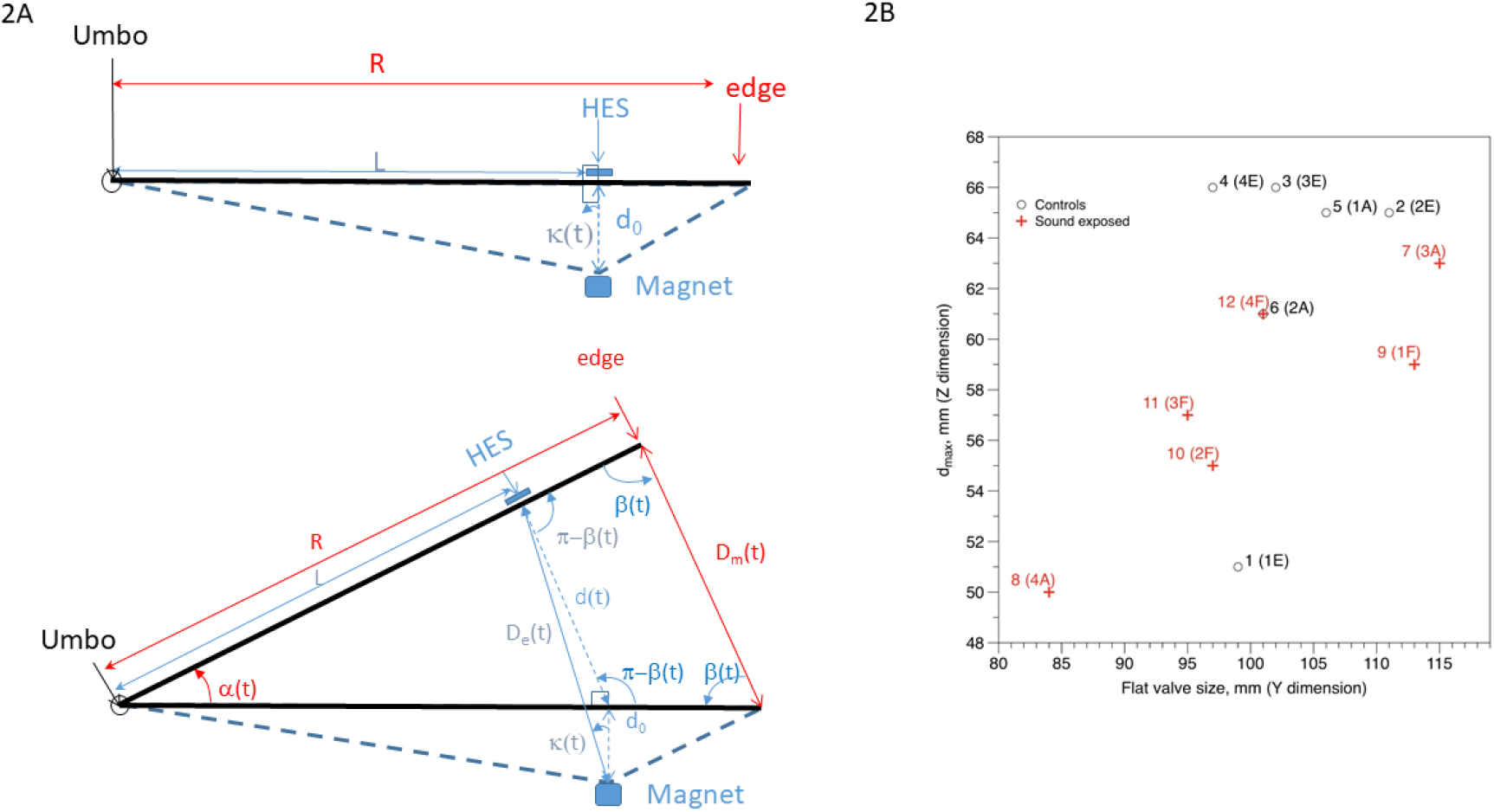
Panels: (A) Geometric representation of a cross-section of a *Pecten maximus* shell showing the point of placement of the Hall-effect sensor (HES) on the flat upper valve and the magnet on the curved lower valve. The valve gape distance to be estimated from the HES voltage readings is *D_m_*(*t*), but the distance that is actually estimated is called *D_e_*(*t*), which is equal to *d*_0_ when the shell is closed. These constraints are taken into account in the developed calibration methods. (B) Range of flat valve sizes and their corresponding maximum intervalve distance for all twelve individuals used in the experiments. The valve sizes depicted here are typical for 3 year old individuals (adults) in the Bay of Brest (France).

The geometry of the shell-HES dynamic system is complex (Figure 2). The valve gape distance to be estimated is the distance *D_m_*(*t*) (in millimeters) between two points located at the edge of each valve; estimates of *D_m_*(*t*) must take into account the relative positions of these two points regarding the relative positions of the HES and the magnet, leading to introducing the distance function, *d*(*t*) (Figures 2A; Equation 3, Table 1). A calibration curve is fitted to a set of *i* = 1,*N* data pair samples *d_i_, U_H_*(*d_i_*) measured on each individual scallop. In the context of the regression, *d_i_* is the explanatory variable and *U_H_*(*d_i_*) is the explained variable. To evaluate the effect of shell opening geometry, we have considered two formulations:

1. The first formulation (Model 1) does not consider the rotation of valves around the hinge. The calibration curve (Table 1, Equation 5) describes the relationship between the unitless sensor output values *U_H_* and a particular distance *d*(*t*) measured at time *t*. It includes 2 parameters *θ* = {*θ*_1_, *θ*_2_} that requires to be estimated. The distance *d*(*t*) is defined as *d*(*t*) = *D_m_*(*t*)*L/R*.
2. The second formulation (Model 2) takes into account the rotation of the valves around the hinge. It introduces two angles *κ*(*t*) and *β* (*t*) (Table 1: Equations 7 and 8), but rests on the similar vector of parameters to estimate, *θ* = {*θ*_1_, *θ*_2_}.

For both formulations, *θ*_1_ is a parameter that is theoretically a function of *I_E_* and *K_H_*. However it also depends on the setup of the measurements (*i.e*. the setup of the data recorder device). *θ*_2_ is theoretically (*i.e*. if the covariance *Cov*(*θ*_1_, *θ*_2_) is equal to zero) a measurement of *d*_0_ (Table 1, Equation 4). These two estimated parameters are linked by the expression 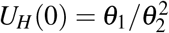.

### 2.2 Inversing calibration functions to build time series of valve gape distances

For the first calibration model, the inverse calibration function (Table 1, Equation 6) had a unique straightforward solution once 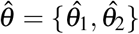 was identified. This was not the case for the second model, which required solving Equation 9 (Table 1) to determine *d*(*t*). This consisted in finding the solutions to a twelve-degree polynomial function:

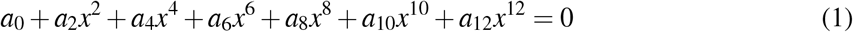

with,

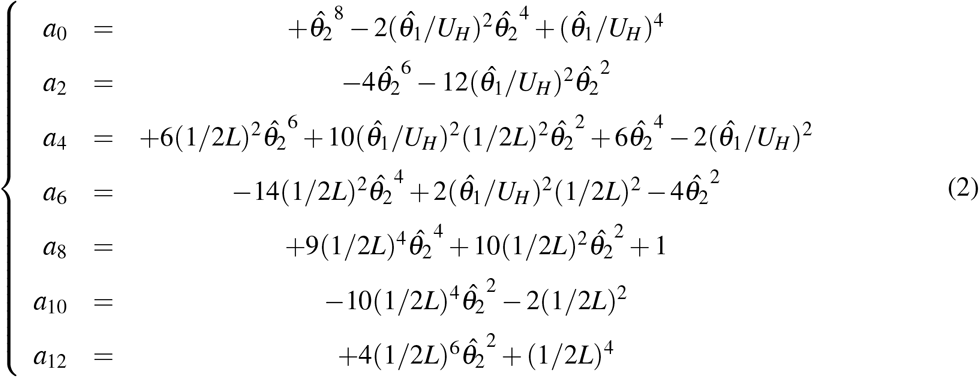

The resolution of equation (1) requires formulating a proposition that, with 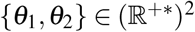, estimated by 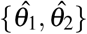 from 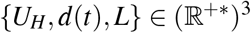, the polynomial function exhibit at least one positive real root, which provides the solution of Equation 9 (Table 1). This proposition was first supported by the fact that inter-valve distances exist (i.e., geometrically, they are described by a real and positive number). Second, this proposition is verified by *a*_12_ > 0. Because non-null coefficients are pairs (hence, −*a*_1_1/*a*_1_2 = 0), roots exhibits positive and negative values, which in turn exhibit a sum and a product equal to zero. These conditions are sufficient to ensure that there is at least one positive real number among all calculated root solutions.

### 2.3 Experimental Data

To illustrate our approach, we used a set of data acquired in experimental controlled conditions. Specimens of *Pecten maximus* of similar sizes (length = 10.2 cm ± 0.9 (SD), n = 12) were hand-collected from the Bay of Brest (France) at a depth of 10 - 15 m in April 2018. Animals were handled with care and their health checked daily. Epibionts were scrubbed gently from shells and scallops were transferred to acclimation conditions for two weeks. They were placed in glass tanks (60 x 50 x 35 *cm*, 105 L), continuously flowed by filtered seawater. The bottom of each tank was covered with a 5 cm deep layer of clean sand and vertical walls were covered with black sheets to prevent visual disturbances (Thomas and Gruffydd, 1971). Lighting in the room was set to a 12h/12h light-dark cycle. Scallops were fed daily with a culture of *Isochrisis galbana* (suspension of 17 10^9^ *cells.L*^−1^).

Experiments started on 16 April 2018 and ended on 22 May 2018. Scallops were removed from their acclimation tanks. A HES (InSb, HW-300A, Asahi Kasei, Japan), cabled and sealed in epoxy resin, was glued to the outer surface of the flat valve at the maximum possible distance from the hinge (Figure 1A). A neodymium magnet (4.8-mm diameter, 0.8-mm thick) was glued on the curved valve surface directly below the sensor. The size of the shell and the linear distance between the sensor and the umbo were measured. The Hall voltage was recorded in dimensionless units with a dynamic strain recorder (DC204R, Tokyo Sokki Kenkyujo Company, Ltd) with a time step of 0.1 second. Four sensors (one per individual) were connected to the same recorder. Neodymium magnets are adjusted to ensure that the Hall voltage reading range was maximal when valves were closed. Experiments were conducted in interconnected plastic tanks (72.5 x 62 x 42 *cm*) filled with seawater up to a water volume of 143 L. The upstream tank was flowed continuously (at 30 *L.h*^−1^) with filtered aerated seawater. The suspension of *I. galbana* was added during daylight conditions (08:00 – 20:00 h) at 1.00 *L.h*^−1^ using a peristaltic pump. The overflow from the upstream tank filled two experimental tanks, each of them splitted in six identical compartments with perforated plastic walls, to minimize interactions between adjacent individuals. Their bottom surface was covered by a 5 *cm* layer of clean sand. They were tented with dark plastic sheets to minimize visual disturbances. Photoperiod was maintained at 12h /12h light-dark cycle using LED light strips (B0187LXUS2, *T_c_* 4500 *K*). Over-flowing water was collected by a downstream tank before being recycled through the system. Two multi-parameter probes (6920 V2, YSI, USA) were placed in the upstream and downstream tanks. Water temperature was equal to 15.89 °C ± 0.89 (SD) in the upstream tank and slightly warmer, 16.17 °C ± 0.69 (SD), in the downstream tank. The salinity was recorded in the upstream tank at 34.02 ± 0.08 (SD).

### 2.4 Data analysis

Calibration measurements were performed for each individual after the experimental observations were completed; the adductor muscles were severed and fourteen glass wedges were successively inserted between the two valves, providing a set of sixteen inter-valve distance values:

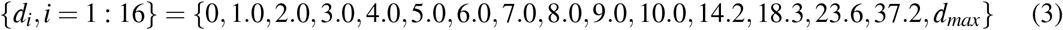

The first value (*d*_1_ = 0) corresponds to a closed shell. The last value (*d_max_*) corresponded to the maximum distance (measured after the adductor muscle was severed), which varied among tested scallops (Figure 2B). The parameter vectors, *θ* = {*θ*_1_, *θ*_2_}, were estimated for Models 1 and 2 by minimizing a weighted least square criterion, *Y_WLS_* (Table 1, Equation 10), using a direct search algorithm (simplex, Nelder and Mead, 1965). The variance-covariance matrix was computed by resampling 100 times the 16 centered residuals with replacement according to the bootstrap method (Efron, 1979) applied to non-linear regression (Efron, 1988).

From estimated parameters *θ* = {*θ*_1_, *θ*_2_}, individual time series of recorded *U_H_*(*d*(*t*)) values were converted in the inter-valve opening distances, *d*(*t*). Time-varying variance was computed using the 100 bootstrap estimates of *θ*. Finally, the first and second derivatives were computed, expressing respectively the speed and acceleration of the distance changes. All calculations were performed using the open source software Scilab (v. 6.0.1, Scilab Enterprise, http://www.scilab.org) and resulting data series were processes with DataGraph (v. 4.3, D. Adalsteinsson, Visual Data Tools Inc., http://www.visualdatatools.com/).

Two additional scallops (y-axis valve lengths equal to 8.1 and 11.5 cm), not equipped with HES, were maintained in acclimation conditions. Their valve movements were annotated from video recordings (GoPro HERO3) over six consecutive days (total time 11 h). Behaviours were categorized according to studies performed on different Pecten species (Braid, 1956; Thomas and Gruffydd, 1971; Wilkens, 1981; Brand, 1991; Robson et al., 2012; Shumway and Parsons, 2016).

## 3 Results

### 3.1 Calibration of valve opening distances

Table 2 presents estimates of the parameters vectors 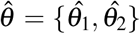 for the two models (M.1, without and M.2, with rotation) for the 12 different individuals. Figure 3 (upper plot) shows an example of a calibration performed for two individuals chosen randomly (Specimens 4 and 9). Estimates of *θ*_1_ and *θ*_2_ are strongly correlated 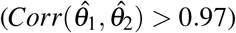. The calculation of *U_H_*(*d*_0_) (dimensionless ordinates at origin) exceeded the maximum value (86,500) for Scallop 1 and 7, for both Models 1 and 2. Overall, ordinates at origin were overestimated for Specimens 1, 2 and 7, and underestimated for Specimen 3, 4, 5, 6, 8, 9, 10, 11 and 12. Values for 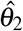 varied from 5.42 ± 0.48 (SE) (Specimen 6) to 10.98 ± 0.75 (SE) (Specimen 11) for Model 1 and from 5.38 ± 0.51 (SE) (Specimen 6) to 10.74 ± 0.78 (SE) (Specimen 11) for Model 2, which is consistent with possible inter-valve distances between the magnet’s position and the edge of the curved valve. Estimated values for 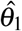 varied between ca. 2.3 10^6^± 0.3 10^6^ (SE) (Specimen 6) and 6.9 10^6^± 0.5 10^6^ (SE) (Specimen 4), and ca. 2.3 10^6^± 0.1 10^6^ (SE) and 6.7 10^6^± 0.4 10^6^ (SE), for Model 1 and Model 2 respectively.

**Table 2.** Model 1 and 2 fitted parameters for the twelve specimens. Best values (estimated from original datasets), Mean and Standard Error values (estimated from 200 bootstrap pseudo-replicates) are represented for 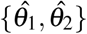 and *U_H_*(*d*_0_), with 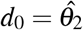 see Table 1. The correlations between the two estimated parameters are represented in 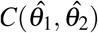, and the last column is for the criteria value (weighted sum of distances between the original data and the function fitted); the best fit between Model 1 and 2 was highlighted in grey for the 12 individuals. The curves shown in Figure 4 are represented using the ‘best’ values of the parameters.

| Specimen | $\hat{\theta}_1$ | $\bar{\theta}_1$ | SE($\hat{\theta}_1$) | $\hat{\theta}_2$ | $\bar{\theta}_2$ | SE($\hat{\theta}_2$) | $U_H(\hat{\theta}_2)$ | $U_H(\bar{\theta}_2)$ | SE($U_H(\hat{\theta}_2)$) | $C(\hat{\theta}_1, \hat{\theta}_2)$ | Criteria |
| --- | --- | --- | --- | --- | --- | --- | --- | --- | --- | --- | --- |
| M.1 |  |  |  |  |  |  |  |  |  |  |  |
| 1 | 3411449 | 3488117 | 759119 | 6.20 | 6.27 | 0.95 | 88598 | 89134 | 8379 | 0.98 | 2400616 |
| 2 | 4323281 | 4330730 | 498246 | 7.45 | 7.45 | 0.60 | 77701 | 77971 | 4010 | 0.98 | 1207225 |
| 3 | 4655724 | 4723360 | 489530 | 8.25 | 8.31 | 0.59 | 68340 | 68328 | 2865 | 0.99 | 355135 |
| 4 | 6821724 | 6908853 | 497360 | 9.67 | 9.74 | 0.48 | 72850 | 72731 | 2220 | 0.99 | 985662 |
| 5 | 3486284 | 3477031 | 224528 | 7.47 | 7.46 | 0.34 | 62390 | 62503 | 1816 | 0.98 | 265508 |
| 6 | 2308542 | 2325220 | 292824 | 5.40 | 5.42 | 0.48 | 79123 | 79317 | 4654 | 0.98 | 692771 |
| 7 | 2842947 | 2897748 | 430503 | 5.73 | 5.80 | 0.60 | 86517 | 86105 | 5496 | 0.98 | 1255806 |
| 8 | 3480304 | 3516563 | 257568 | 7.16 | 7.21 | 0.38 | 67737 | 67588 | 2439 | 0.98 | 205485 |
| 9 | 5991495 | 6008167 | 600860 | 10.41 | 10.42 | 0.68 | 55189 | 55270 | 1971 | 0.99 | 203447 |
| 10 | 4851406 | 4882129 | 270263 | 9.91 | 9.94 | 0.37 | 49350 | 49344 | 1138 | 0.98 | 674078 |
| 11 | 5328872 | 5442886 | 545266 | 10.83 | 10.98 | 0.75 | 45371 | 45112 | 1841 | 0.98 | 970991 |
| 12 | 2889610 | 2902979 | 185142 | 7.23 | 7.26 | 0.33 | 55128 | 55081 | 1651 | 0.98 | 253263 |
| M.2 |  |  |  |  |  |  |  |  |  |  |  |
| 1 | 3395297 | 3536801 | 926412 | 6.18 | 6.30 | 1.13 | 88771 | 89406 | 9500 | 0.98 | 2446105 |
| 2 | 4301616 | 4325321 | 528357 | 7.43 | 7.44 | 0.63 | 77860 | 78239 | 4284 | 0.98 | 1250001 |
| 3 | 4627845 | 4603136 | 526754 | 8.21 | 8.17 | 0.65 | 68498 | 68883 | 3269 | 0.99 | 465945 |
| 4 | 6780506 | 6787939 | 468733 | 9.63 | 9.63 | 0.45 | 73006 | 73076 | 2058 | 0.98 | 904649 |
| 5 | 3472454 | 3493657 | 276089 | 7.45 | 7.47 | 0.41 | 62487 | 62613 | 2197 | 0.99 | 254702 |
| 6 | 2299438 | 2305817 | 306529 | 5.38 | 5.38 | 0.51 | 79264 | 79785 | 4871 | 0.98 | 715674 |
| 7 | 2833806 | 2869798 | 378464 | 5.71 | 5.76 | 0.54 | 86637 | 86629 | 5458 | 0.98 | 1281171 |
| 8 | 3457813 | 3496710 | 252566 | 7.13 | 7.17 | 0.37 | 67914 | 67952 | 2313 | 0.98 | 202756 |
| 9 | 5955095 | 6072554 | 619694 | 10.37 | 10.50 | 0.72 | 55308 | 55027 | 2133 | 0.99 | 201116 |
| 10 | 4820499 | 4828595 | 280198 | 9.87 | 9.87 | 0.40 | 49460 | 49512 | 1225 | 0.99 | 629871 |
| 11 | 5293625 | 5281907 | 567485 | 10.78 | 10.74 | 0.78 | 45473 | 45756 | 1938 | 0.99 | 914811 |
| 12 | 2878374 | 2897727 | 166986 | 7.22 | 7.25 | 0.30 | 55215 | 55095 | 1540 | 0.98 | 238846 |

**Figure 3:**
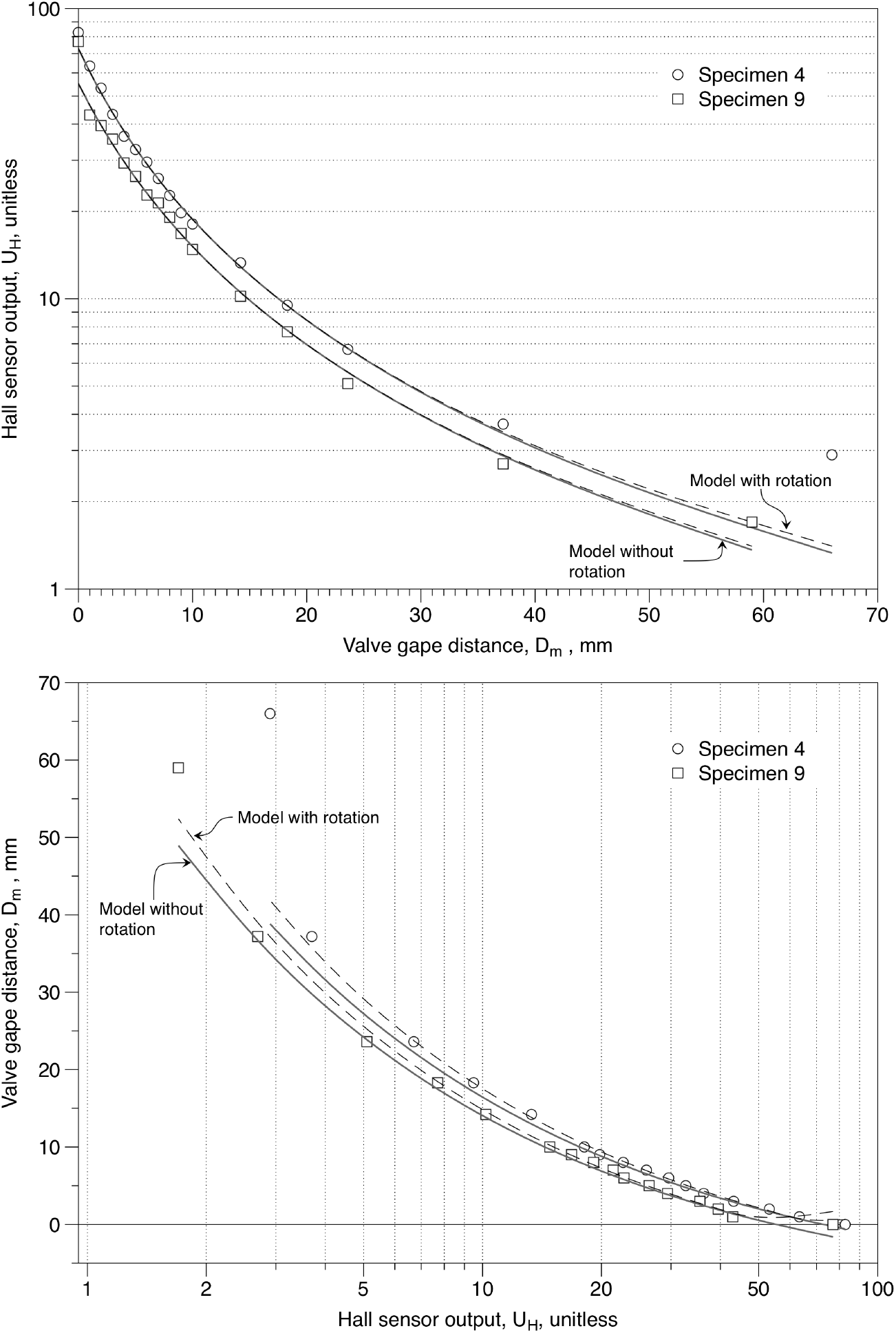
Example of calibration curves estimated for two specimen 4 and 9 (randomly chosen from the 12 specimens). Fit parameters are provided for all specimens in Table 3. Empty symbols represent observations and lines correspond to estimated calibration curves, calculated with the optimized parameters. The solid line represent the curve calculated from the model without taking into account rotation around the hinge (Model 1) and the dash line correspond to the curve calculated with the model that takes into account rotation (Model 2). The upper figure shows the direct optimized curves, where the distances *D_m_* were controlled and the voltage-related values *U_H_* were measured, whereas the lower part shows the inversed (reciprocal) curve allowing one to calculate variations of *D_m_* from a time series of measured *U_H_*.

We did not detect any bias in any of the estimates. Parameter estimates were similar for Model 1 and Model 2. The criterion value (sum of square weighted differences between data and model estimates) was lower for Model 1 than for Model 2 in five cases (Specimens 1, 2, 3, 6 and 7), and was lower for Model 2 than Model 1 for the other seven specimens. Figure 3, upper graph, shows that the Model 2 (with rotation around the hinge) produced, for both specimens, curves (dash line) that fit better to the highest distance values when the Hall voltage decreased and the angle between the magnet and the sensor axes increased.

The model 2 that takes into account rotation of valves around the hinge is assumed to provide unbiased estimators, in contrast to the model 1 that does not account for rotation. Thus, for small Hall voltages (due to large distance between the HES and the magnet), a small discrepancy between the two models induces a large difference between the corresponding estimated gape distances (Figure 3, lower graph), which shows that rotation is important to these estimates. This means that when time series of gape distances are reconstructed from valvometry data, the gape distances estimated by model 1 will be significantly biased whenever rotation has to be accounted for (Figure 4).

**Figure 4:**
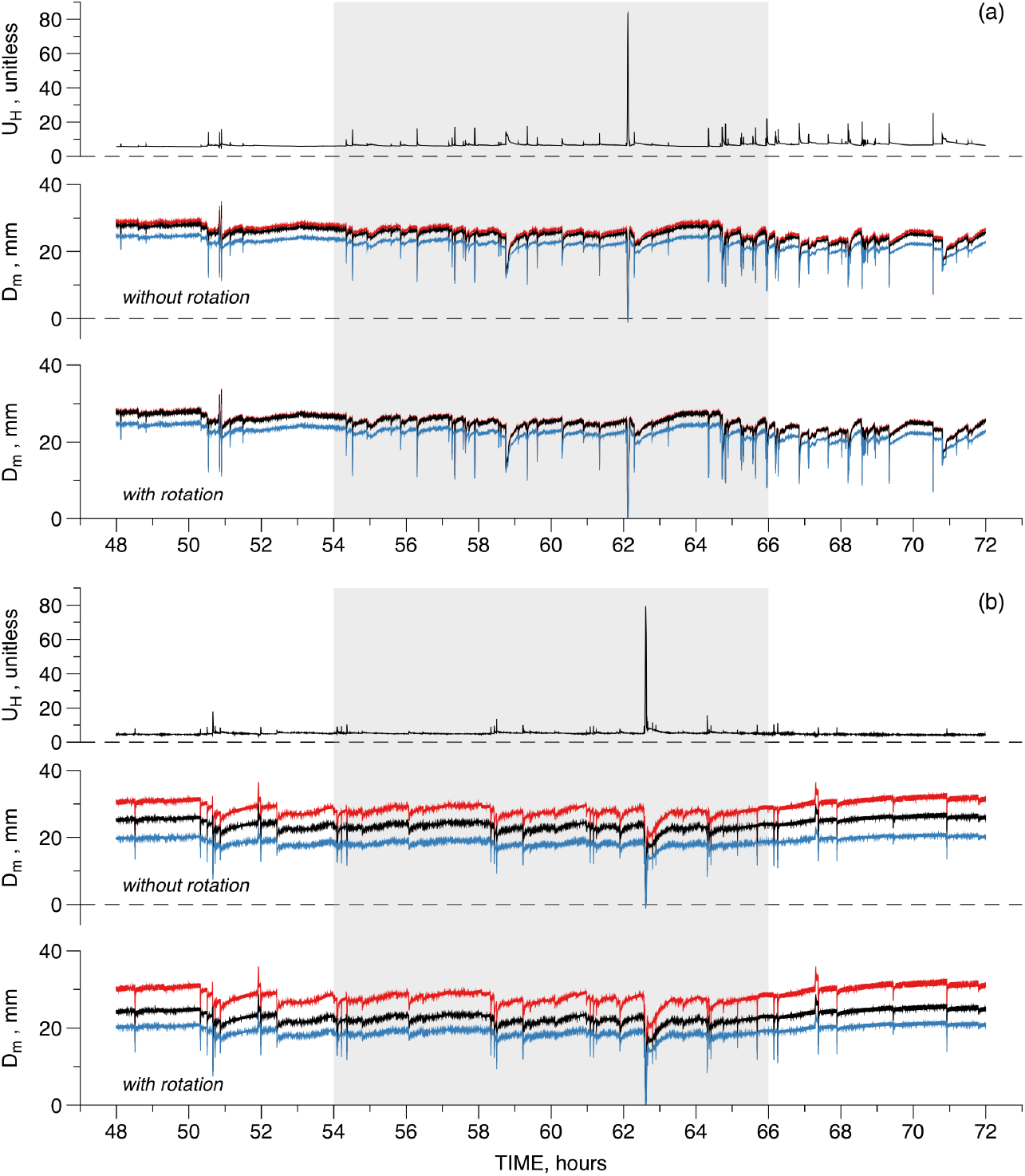
Example of the conversion of Hall voltage records (upper graph) to the time series of opening distances calculated from calibrations of Model 1 (middle graphs) and Model 2 (lower graphs) for Specimens 4 (a) and 9 (b) over a 24-h period. Calibrated parameters and related Distances, *D_m_*, variability was estimated with 100 bootstrap replicates. The two lines surrounding the best estimates of the time series, after inversion of the calibration curves, representing time series of standard errors, illustrate the resulting variability. The white bands indicate daytime (12 h of light, between 14:00 and 20:00, and between 8:00 to 14:00), whereas the grey band indicates night times (12 h of darkness, between 20:00 and 8:00).

For high Hall voltages (when the HES nears the magnet), the discrepancy between the two models is small. However, the lower graph of Figure 3 shows that the inversed function of model 1 (no rotation of the valves around the hinge) allows distance estimates to become negative. On the contrary, model 2 that accounts for rotations, inflects values in the proximity of zero, preventing estimated distances from becoming negative.

### 3.2 Converting time series of Hall voltage measurements to valve gape distances

Figure 4 represents the time series measured and calculated for inter-valve distances estimated for Specimens 4 and 9 over a 24-h period. The calculation of inter-valve distances, performed with inversion of Model 1 (middle graph) and Model 2 (lower graph), included the variability around the best estimates, using the bootstrap replicates generated for the calibration.

Table 3 provides results for all 12 series. The average inter-valve distance estimates is similar for the two models. The average inter-valve distance was ca. 27 mm but with a minimum of ca. 18 mm for Specimen 10 and maximum of ca. 44 mm for Specimen 12. This range represents an inter-valve distance relative to *d_max_* ranging from 30% to 73% (average 46% for Models 1 and 2). The uncertainty (in terms of SE) around the best estimate varied from 3 to 14 mm and was lower for Model 2 than for Model 1 except for Specimens 5 and 12. The average uncertainty (in terms of SE) was about 7 mm for Model 1 and 6 mm for Model 2. There was no strong relationship between the average inter-valve distance and the estimated variability. Figure 4 shows that the uncertainty distribution around the best estimate varies from one individual to the other, with varying skewness.

**Table 3.** Performance of Models 1 and 2 estimating variations of inter-valve distances recorded with HES sensors on 12 individual specimens. The bootstrap replicates calculated for the calibrations were used to estimate the mean *M*(Δ) and the standard error, *SE* (Δ), of the dispersion around the average inter-valve distances.

| M.1 |  |  |  |  |  | M.2 |  |  |  |  |
| --- | --- | --- | --- | --- | --- | --- | --- | --- | --- | --- |
| Specimen | $\bar{D}_m(t)$ | $\min(D_m(t))$ | $\max(D_m(t))$ | $\bar{\Delta}$ | $SE(\Delta)$ | $\bar{D}_m(t)$ | $\min(D_m(t))$ | $\max(D_m(t))$ | $\bar{\Delta}$ | $SE(\Delta)$ |
| 1 | 26.55 | 3.73 | 39.68 | 14.24 | 2.56 | 26.65 | 3.73 | 40.07 | 10.56 | 1.96 |
| 2 | 19.80 | 6.44 | 41.76 | 4.26 | 0.65 | 19.81 | 6.43 | 42.17 | 4.10 | 0.67 |
| 3 | 29.13 | 4.70 | 49.07 | 5.56 | 0.62 | 29.23 | 4.70 | 49.74 | 5.07 | 0.58 |
| 4 | 23.94 | -0.66 | 32.10 | 3.70 | 0.38 | 23.99 | 0.00 | 32.33 | 2.74 | 0.28 |
| 5 | 27.33 | 0.06 | 71.78 | 10.35 | 1.98 | 27.45 | 0.07 | 74.64 | 14.11 | 2.65 |
| 6 | 27.59 | 2.95 | 70.61 | 8.05 | 1.03 | 27.68 | 2.95 | 73.11 | 6.83 | 0.87 |
| 7 | 31.01 | 2.91 | 45.98 | 9.26 | 1.34 | 31.15 | 2.92 | 46.52 | 4.74 | 0.69 |
| 8 | 31.54 | 8.67 | 58.01 | 5.43 | 1.00 | 31.68 | 8.66 | 59.04 | 4.15 | 0.79 |
| 9 | 21.84 | 0.38 | 27.43 | 9.13 | 0.83 | 21.88 | 0.39 | 27.54 | 7.06 | 0.62 |
| 10 | 17.99 | 3.27 | 25.10 | 4.21 | 0.68 | 18.00 | 3.28 | 25.15 | 3.91 | 0.66 |
| 11 | 24.51 | -1.03 | 37.98 | 4.81 | 1.41 | 24.57 | 0.00 | 38.26 | 3.88 | 1.14 |
| 12 | 44.27 | 19.78 | 51.46 | 6.21 | 0.41 | 44.70 | 19.79 | 60.57 | 6.26 | 0.44 |

### 3.3 Inferring the valve-gape dynamics

Figure 5 shows the variations in inter-valve distance (upper graph), movement velocity (middle graph), and acceleration (lower graph), using the calibration function that took into account rotation around the hinge axis (model 2), for Specimens 4 and 9.

**Figure 5:**
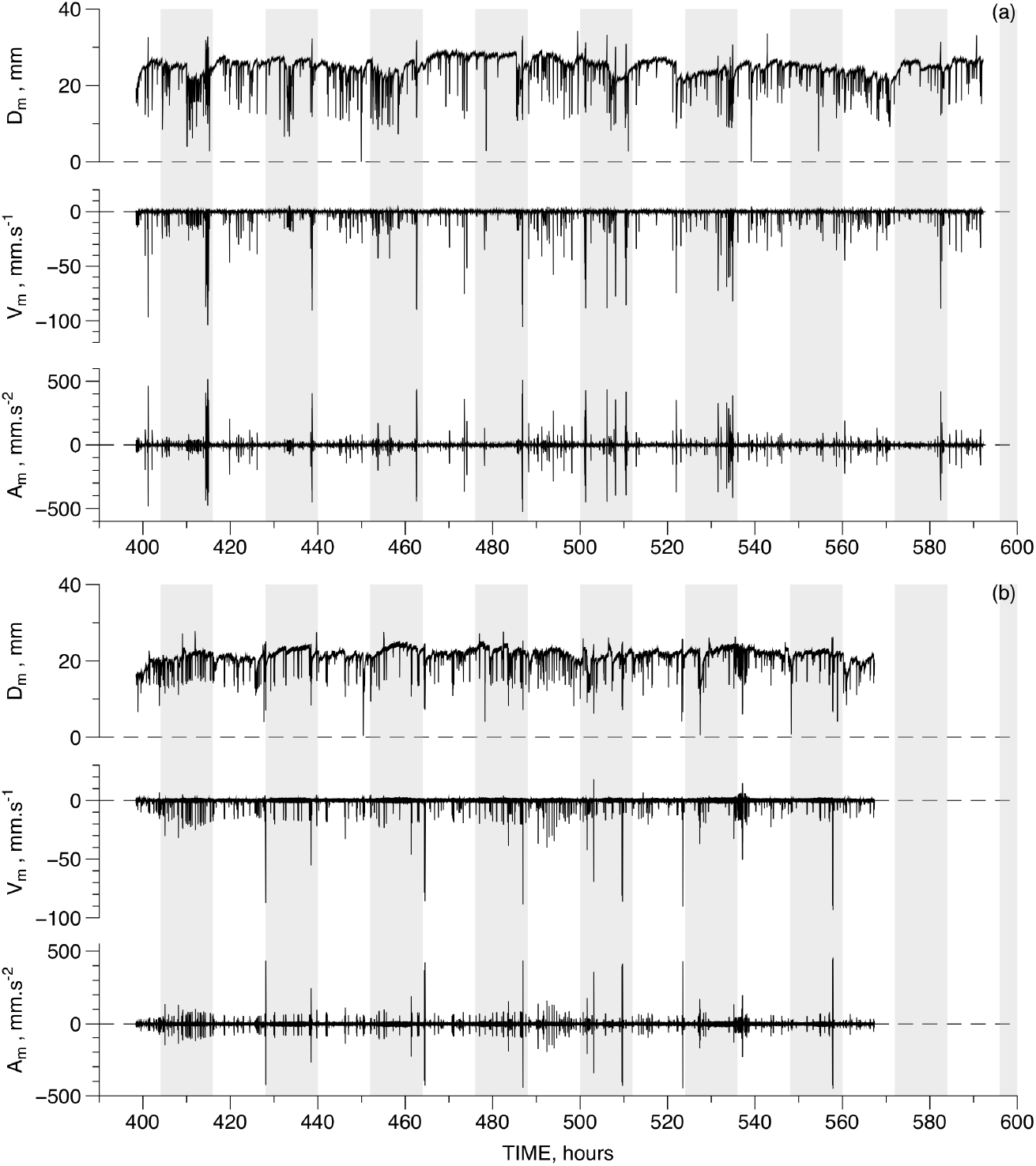
Example of valvometry time series’ recorded for about 8 d for Specimens 4 (a) and 9 (b). Time series were constructed from the calibration of Model 2. In both cases (a) and (b), the upper plot shows the estimated inter-valve distances over time, the middle plot shows the calculated velocity of valve movement over time (positive values indicate opening and negative values indicate closing), and the lower plot shows the calculated acceleration over time. The white bands indicate daytime (12 h of light, between 8:00 and 20:00), whereas the grey bands indicate night times (12 h of darkness, between 20:00 and 8:00).

Table 4 shows results for all 12 specimens. Time series were recorded for 194 h (Specimen 1 to 4) and 169 h (Specimen 5 to 12) respectively, except for Specimen 10 (160 hours, because the first 9 h were not exploitable). The averages speeds and accelerations were equal to zero for all the 12 specimens. Some individual dynamics exhibited sharp bursts of speed approaching 200 *mm.s^−1^* and accelerations of up to ca. 3000 *mm.s*^−2^). Considering that the order of magnitude of distance estimated during the experiments was 20 mm, these values suggest that limits of speed and accelerations for the 10 Hz measurement frequency were reached. These peaks can thus be considered as discrete events in the continuous dynamics.

**Table 4.** Estimates of speed, acceleration and number of peaks using calibration Model that takes into account rotation around the hinge. The data analyses were performed for 12 specimen. The time series were recorded at 10 Hz (1 value every 0.1s). Averages of speeds (v, *mm.s*^−1^) and accelerations (a, *mm.s*^−2^) were equal to zero (with a 2 digits precision), even if the maximum closing speed (*v_down_*) is usually higher than the maximum opening speed (*v_up_*). There were discrepancies in the related accelerations but the maximum acceleration during closing event (*a_down_*) were very similar to the maximum acceleration during valve opening (*a_up_*). The two last columns shows the number of peaks estimated and the related frequency (number of peaks divided by the duration in hours).

| Specimen | Dur. (h) | Dur. (d) | $\bar{v}$ | $v_{down}$ | $v_{up}$ | $\bar{a}$ | $a_{down}$ | $a_{up}$ | nb peaks | Freq. |
| --- | --- | --- | --- | --- | --- | --- | --- | --- | --- | --- |
| 1 | 194 | 8.07 | 0.00 | -200.10 | 75.55 | 0.00 | -1366.02 | 846.11 | 620 | 3.20 |
| 2 | 194 | 8.07 | 0.00 | -218.26 | 13.43 | 0.00 | -1123.70 | 1053.69 | 408 | 2.11 |
| 3 | 194 | 8.07 | 0.00 | -194.02 | 192.81 | 0.00 | -1923.99 | 1934.14 | 561 | 2.90 |
| 4 | 194 | 8.07 | 0.00 | -105.71 | 5.83 | 0.00 | -527.52 | 517.73 | 413 | 2.13 |
| 5 | 169 | 7.03 | 0.00 | -323.65 | 313.98 | 0.00 | -3188.19 | 2812.54 | 605 | 3.58 |
| 6 | 169 | 7.03 | 0.00 | -314.06 | 314.87 | 0.00 | -3144.64 | 2327.31 | 2189 | 12.97 |
| 7 | 169 | 7.03 | 0.00 | -246.14 | 182.59 | 0.00 | -1820.43 | 1393.26 | 7862 | 46.59 |
| 8 | 169 | 7.03 | 0.00 | -210.44 | 43.63 | 0.00 | -1032.49 | 1027.37 | 11273 | 66.80 |
| 9 | 169 | 7.04 | 0.00 | -93.39 | 18.16 | 0.00 | -451.41 | 456.45 | 421 | 2.49 |
| 10 | 160 | 6.67 | 0.00 | -79.17 | 5.35 | 0.00 | -404.97 | 390.26 | 208 | 1.30 |
| 11 | 169 | 7.04 | 0.00 | -123.57 | 9.77 | 0.00 | -629.57 | 605.95 | 524 | 3.10 |
| 12 | 169 | 7.04 | 0.00 | -229.80 | 215.92 | 0.00 | -2132.79 | 1982.17 | 231 | 1.37 |

On average, 440 peaks were estimated from the 12 series (between 208 peaks for Specimen 10 and 620 peaks for Specimen 1). Therefore, there was *ca*. 1 peak every 25 min on average (from every 15 min for Specimen 1 to 45 min for Specimen 10). Most of the peaks (about 95% in average) were single closing events. The speed values distributions were strongly asymmetrical, with higher speeds for valve closings than for valve openings. A few exceptions occurred when a shell inter-valve distance oscillated quickly creating a symmetrical distribution. In contrast, distributions of acceleration values were always close to symmetric.

### 3.4 Behaviour

Seven distinct behaviours were identified from video recordings (Table 5). They were categorized as: (1) a single contraction followed by slow relaxation (related to partial or complete closure) or (2) repeatedly rapid contractions (related to jumping, rotation, burying, and swimming). Of these behaviours, partial closure (1, Table 5) was the most common. It is characterized by a rapid closing movement, followed by a period of slow re-opening, which can last over one hour. It is similar to total closure (2, Table 5), which implies the complete retraction of the mantle rather than a partial one observed in behaviour 1. Jumping and rotating behaviours resemble each other in the valvometry series; they began with a strong valve opening, followed by strong and rapid contractions that expelled water contained in the pallial cavity, thus provoking the animal to either jump backwards or rotate swiftly. Swimming and burying behaviours (3 and 6, Table 5) were characterized by series of contractions, rapid and frequent when swimming, or rapid and irregularly spaced when digging into the tank sediment. After digging, the scallop would usually continued to expel sand particles *via* a series of slow frequency, large amplitude movements.

**Table 5.** Six movement behaviours associated with valve openings observed in video recordings. Descriptions of behaviours by Thomas and Gruffydd (1971), Wilkens (1981), Brand (1991), and Robson et al. (2012). Movements described as ‘rapid’ or ‘sudden’ indicate sharp changes in inter-valve distance (d) at 0.1 sec intervals (the time-step used in valve-gape experiments). It is important to recall here that these experiments were not designed to test any proximal or distal causes for these movements.

| Behaviour | Description |
| --- | --- |
| 1-Partial Closure | Valve suddenly closes partially then re-opens slowly. It occurs when environmental conditions change (*e.g.* change of light, sudden movements of another individual). It may be accompanied by feces expulsion. |
| 2-Total Closure | Valves close quickly, for few seconds, then reopen slowly over several minutes and up to an hour. It may be accompanied by feces expulsion. |
| 3-Swimming | Rapid succession of valve opening and closing movements inducing a forward movement, off the substrate. |
| 4-Jumping | Valve movements start by a large valve opening movement followed by a strong, rapid contraction expelling water in the pallial cavity and causing the animal to ‘jump’ backwards. |
| 5-Rotation | Valve movements resemble to jumps but individuals rotate while remaining on the substrate. |
| 6-Burial/Digging | Relatively long (several minutes) series of irregularly spaced rapid openings and closures, which force the shell into the sand at the bottom of the tanks. It may be followed by lower frequency movements which create a depression in the sand around the individual. |

## 4 Discussion

### 4.1 Technical limitations of valvometry for characterising valve opening

A Hall-Effect sensor, as any electronic device, has intrinsic limitations. It produces electrical noise expressed as a random voltage fluctuations (Ejsing, 2006). Several potential sources exist, including: the transducer itself, the signal amplifier, or the environment in which it is placed (Ramsden, 2011). The electronic variability generated by Hall transducers (Ejsing, 2006) have been characterized as Johnson-Nyquist noise (due to the thermal-induced agitation of the charge carriers in the semi-conductor) and pink noise (noise with a power spectral density function of the inverse of its frequency). Both types of noises have been reported in Hall-effect sensor valvometry (Bril, 2010).

The second type of limitations on valvometry recordings is induced by the shell morphology of the individual specimens used. As for many other biological measurements (Schwamborn, 2018; Marshall and White, 2019), valvometry only provides a one-dimensional estimate of a three-dimensional process. In this study, we accounted for only two dimensions; this requires that method protocols include two precautions: first is to ensure that the sensor and the magnet are aligned properly across from each other during installation, and secondly that any changes in valve gape distances measured at the edge of a shell are homothetic. Homothetic measurements in this case mean that both halves of the sensor are placed on shell valves in the same cross-sectional plane (on Figure 1A, this would by the Y-Z plane). Homotheticity is rarely mentioned in present-day works, but was discussed in very early valvometry studies (*e.g*. Marceau, 1906, 1909) where three-dimensional discrepancies in shell movements were noted in numerous different bivalve species, including pectinids. Marceau (1909, p. 458), characterized it as a *”mouvement de bascule autour de l’axe dorso-ventrale”*, which translates in English as a pitching or swinging motion of the valves around the dorsal-ventral axis. He attributed this to the differential action of the adductor muscle(s), and states that the motion is more pronounced in species with two adductor muscles, than in those with one, like the pectinids. The requirement of homothetic measurements is a fundamental assumption for reconstruction and validation of valve gape time series. When sensitive sensors like HES are used, it is important to address possible sources of asymmetry, whether they concern shell shape or the movement itself. For example, when valvometry is used for long time periods (that is over several months, even years), existing asymmetries can be exacerbated during shell growth. Shape asymmetries can be tested for by modelling shell morphodynamics (D’Arcy Thompson, 1945); for a perfectly- and symmetrically-shaped shell (*i.e*., with equiangular spiral properties), generated by geometric growth, the homothetic condition can be assumed. However, geometric growth is rarely observed in real shells (Guarini et al., 2011). Therefore, homotheticity should be tested by using several valvometers placed at different points on a single individual; the resulting observations can then be compared with homothetic predictions. Because asymmetries and other morphological particularities may affect valve movements differently from one individual specimen to another, it would also be necessary to provide a morphodynamic description for each specimen used in order to detect specific points of interest where unbiased calibrations can be performed. Our results suggest that additional experiments are necessary to explore the importance of these issues using different species and observation periods with higher resolution sensors.

### 4.2 Calibrating Hall-effect sensors for bivalve valvometry

HES has already been employed in a number of valvometry studies involving not only sessile species, such as *Mytilus galloprovincialis* (Comeau et al., 2019) and *Crassostrea virginica* (Clements et al., 2018; Casas et al., 2018 and Comeau et al., 2012), but also mobile species such as scallops (*Pecten maximus*) (Robson et al., 2009) that are capable of swimming and jumping. Many models have been developed for calibrating valve dynamics, for example with polynomial fitting (Robson et al., 2009; Bamber and Westerlund, 2016), linear fitting (Liu et al., 2016) or negative exponential functions (Miller and Dowd, 2017). The models used here (Table 1) differ in that they are mechanistic, based on physical principles related to variations in the magnetic field intensity; it varies with the inverse of the squared distance between the Hall-effect sensor and the magnet, when the Hall-effect sensor is powered at a constant current and intensity. In addition, we incorporated a function that accounts for the valve rotation around the hinge axis, which modifies the relative position (hence the alignment) of the HES and the magnet.

Our work highlights four aspects of valvometry studies that remain poorly characterized:

1. The principle that the magnetic field intensity varies with the inverse of the distance between the magnet and the Hall-effect sensor (HES) is only rigorous when the HES and the magnet are perfectly aligned (Popovic, 2003). Small deviations from this alignment, which are difficult to avoid when gluing a sensor and magnet to shell surfaces with varying topographic textures, can lead to strong fluctuations in the resulting signal.
2. The sensitivity of the calibration functions decreases when distances d increases. It is expressed by differentiating *U_H_*(*d*) with distance, d:

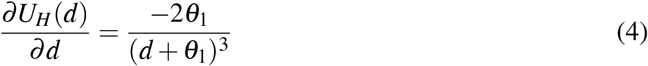

which converges toward zero [faster than *U_H_*(*d*) declines] when d increases. Conversely, the sensitivity of the inverse function is:

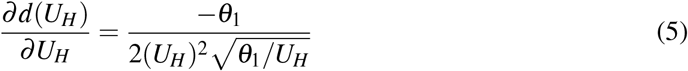

which is maximal when *U_H_* (*d*) is close to zero (distance converging to *d_max_*) and minimal when *U_H_* (*d*) is at its maximum. This problem can be mitigated by positioning the sensor and the magnet closer to the hinge. However, positions close to the hinge would result in a general loss of sensitivity in distance estimates, since uncertainties are amplified by the ratio between the distance from the hinge to the edge divided by the distance from the hinge to the sensor.
3. The calibration function should account for environmental variability, like temperature fluctuations (Liebsch, 2002; Popovic, 2003). The HW-300A HES used in this study has a variation rate to temperature changes equal to −0.018 °C^−1^ between 0 °C to 40 °C. This rate of change is six times greater than that of the majority of transducers, which are closer to 0.003 °C^−1^ (Ramsden, 2011). However, in our case, the temperature effect was relatively small considering that the variations of temperature that we recorded experimentally (not more than 2 °C). It is nevertheless small regarding the usual range of temperature fluctuations that individuals experience in natural environments (about 10 °C to 20 °C), especially for organisms inhabiting intertidal areas (Miller and Dowd, 2017).
4. The calibration model accounting for rotation around the hinge (Model 2) is unbiased, hence should be favoured over the Model 1, which does not take into account rotation. However, in practice, Model 2 only slightly improved fits to our data (see Figure 4), likely because of the variability induced by the relative positions of the HES and magnet. The computational complexity required to invert the curve (to calculate distances from Hall-Effect data) may therefore lead one to prefer Model 1 as a satisfactory approximation, especially when the inter-valve distances remain small (*e.g*., for species like *Mytilus galloprovincialis*, Comeau et al., 2019, or *Crassostrea virginica*, Comeau et al., 2012).

### 4.3 Characterizing the dynamics of inter-valve opening in term of behaviour

Most of the time, scallops open their shells at inter-valve distances that are much lower than the maximum value (on average, 40% of *d_max_*). Therefore, the adductor muscle make a sustained effort to maintain valves in this partially closed position. This continuous effort is ponctuated by discrete events of rapid valve movements, due to two categories of behaviours: single closures, sometimes preceded by a relaxation and followed by a slower re-opening, and a series of flapping behaviours, usually preceded by a sharp relaxation and followed by a slow re-opening. Pectinids, and especially *Pecten maximus*, are known to settle into sediment by nestling quickly when they are displaced or disturbed (Minchin et al., 2000). Once burried in the sediment, single movement events still occur, to expel water, sediment particles, or faeces. They also rotate to orient the valve opening to face the direction of water flow (Shumway and Parsons, 2016). In addition, shell closures and jumps have been described as reactions to external stresses, such as predator contact, vibration, and changes in light and temperature (Thomas and Gruffydd, 1971). Other stress-induced movements described in the literature, were not observed during this present study. In particular, we did not observe flipping (Brand, 1991) or flinching behaviours (Day et al., 2017). In general, valvometry data alone cannot be used to assign one particular movement to a particular behaviour. This would require formulating behavioural and ecophysiological models and performing additional experimentation to link environmental and biological constraints with appropriate responses.

## 5 Conclusion

Hall-Effect Sensors are suitable for valvometry applications, however, to provide reliable unbiased estimators, they require a calibration that takes into account both the physical properties of the sensor and the geometry of the shell. In this article we describe the challenges of constructing time series of the variations of valve gape distance from the inversion of calibration curves. Using pseudo-replicated parameter estimates, we report that the calculated time-varying dispersions and uncertainties of the inter-valve distance are far from being negligible. We have concluded that these problems of variability and uncertainty could be improved by increasing the precision in the positioning of the HES-magnet contraption, for future studies.

Time series data by themselves are insufficient for providing causal explanations about observed patterns. We have concluded only that the dynamics of bivalve shell movements can be described as hybrid, characterized by a continuous convergence to a steady-state opening value interrupted by discrete events of fast movements. This dynamic feature precludes the use of classical time series statistical analyses, including methods designed to infer variability, filter data, or extract trends. Methods to analyse such data are often complicated and require a substantial modelling effort (Yaghoubi and Fainekos, 2018; Collins et al., 2012). The next step will be to combine experimental datasets with numerical models and optimisation techniques. To explore valve gape dynamics from a physiological perspective, the quantification of ‘effort’ would imply:

- expressing speed and acceleration of valve movements in terms of angular momentum;
- designing a hybrid dynamic model of the adductor muscle activity to represent the dynamics of valve movements; and,
- describing explicit forcings that provoke changes in muscle activity.

A first step towards reaching these goals is to convert inter-valve distances into opening angles (Wilson et al., 2005). However, the opening angle, as a geometric approximation of shells, has little biological relevance due to the way the morphospace of mollusc shells is organized. The opening of a shell should be considered in all three dimensions, not as a one-dimensional inter-valve distance or as a two-dimensional plan containing a triangle. We suggest that valvometry patterns will remain incomplete without developing a 3D dynamic model that considers shell morphology plans in a way that relates valvometry measurements more closely with biological processes.

Finally, under stress, Pectinidae, as other bivalves, tend to close their shell to ”shelter”, and this constitutes an increase of physiological effort which could be treated as an ”environmental impact”. Because valvometry can record rapid movements linked to shell openings/closing movements, *in situ*, it has a strong potential as a method for identifying behavioural changes during exposures to environmental perturbations. Nevertheless, the consequences of environmental disturbances on the physiology of bivalves should be properly assessed within a quantifiable, causal framework before drawing any correlative interpretation between behaviour and environmental perturbations.

## Acknowledgements

This text is omitted for the preprint.

## Competing interest(s)

This text is omitted for the preprint.

## Notes

### Competing Interest Statement

The authors have declared no competing interest.

